# Neural circuits for long-range color filling-in

**DOI:** 10.1101/223156

**Authors:** Peggy Gerardin, Clément Abbatecola, Frédéric Devinck, Henry Kennedy, Michel Dojat, Kenneth Knoblauch

**Author notes:** Correspondence should be addressed to: Peggy Gerardin & Kenneth Knoblauch, INSERM U1208 Stem-Cell and Brain Research Institute, 69500 Bron, France.

## Abstract

Surface color appearance depends on both local surface chromaticity and global context. How are these inter-dependencies supported by cortical networks? Combining functional imaging and psychophysics, we examined if color from long-range filling-in engages distinct pathways from responses caused by a field of uniform chromaticity. We find that color from filling-in is best classified and best correlated with appearance by two dorsal areas, V3A and V3B/KO. In contrast, a field of uniform chromaticity is best classified by ventral areas hV4 and LO. Dynamic causal modeling revealed feedback modulation from area V3A to areas V1 and LO for filling-in, contrasting with feedback from LO modulating areas V1 and V3A for a matched uniform chromaticity. These results indicate a dorsal stream role in color filling-in via feedback modulation of area V1 coupled with a cross-stream modulation of ventral areas suggesting that local and contextual influences on color appearance engage distinct neural networks.

## 1. Introduction

Surface color appearance depends on multiple factors. In the absence of contextual information, color appearance largely depends on surface spectral information and will be referred to as *surface-dependent color*. Nevertheless, contextual features influence and modulate color appearance. The contributions of contextual influences to surface color appearance have interested scientists for many years and have been successfully studied via a number of visual phenomena, including simultaneous contrast (Chevreul 1839), neon-color spreading (Varin 1971), and, more recently by the Watercolor Effect (Pinna, Brelstaff, and Spillmann 2001). Surface color perception that is driven by remote contours is referred to as filling-in. The above studies suggest that the color appearance of surfaces depends on combining local surface information with contextual information, such as from distant edges. The latter we refer to as *edge-dependent or induced* color. In contrast to the progress made on the psychophysics of these phenomena, little is known about the distributed neural representation of edge-induced filling-in percepts and precisely to what extent identical percepts induced by edge and surface information depend on common or distinct neural networks.

A common hypothesis is that surface- and edge-dependent percepts generate equivalent neural activity at early visual stages (Komatsu 2006). Ratliff and Sirovich (1978) noted that convolution with center-surround like filters both of step-edges and of the spatial transients that induce filling-in (e.g., Craik-O’Brien-Cornsweet effects) results in nearly identical images (Figure 1). One possibility is that the center-surround receptive field organization in early vision could leads to edge transients and surfaces generating equivalent neural response profiles that the observer would perceive similarly.

**Figure 1:**
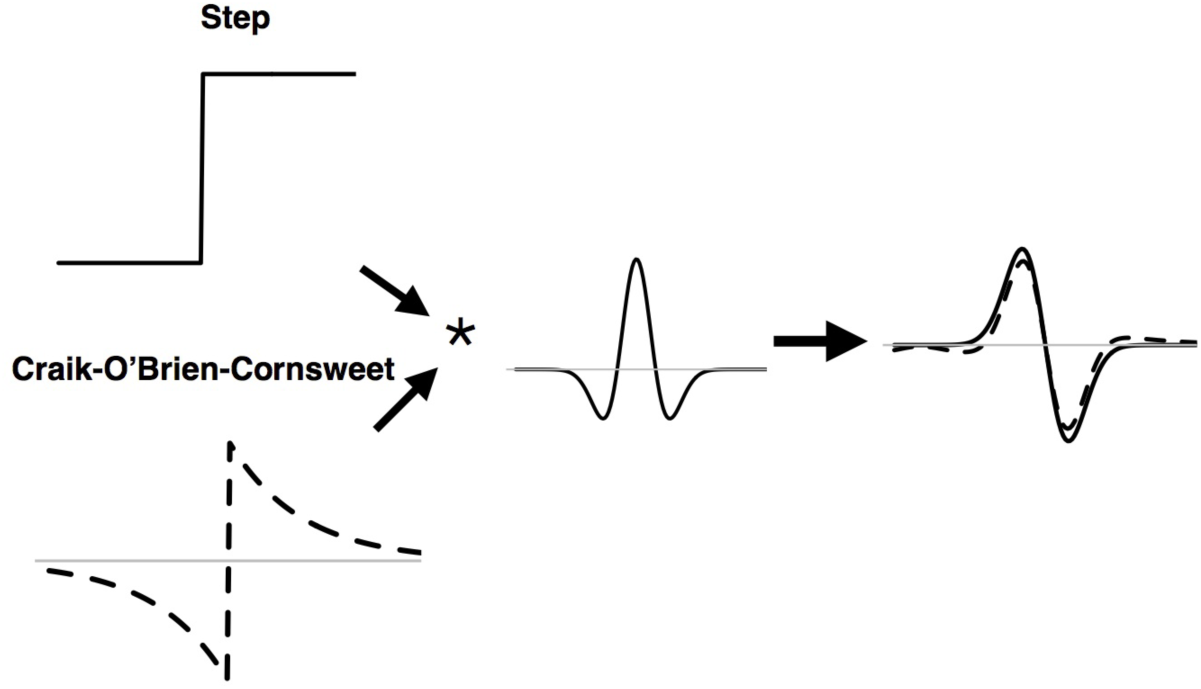
Equivalence of responses for step and gradient intensity profiles under convolution with a center-surround weighting function.

Both a step change in intensity (solid curve, upper left) and the exponential ramp transient for a Craik-O’Brien-Cornsweet effect (dashed curve, lower left) when convolved (indicated by “*”) with a center-surround weighting function (center, the negative second derivative of a Gaussian) yield nearly identical response profiles (right, solid and dashed curves, respectively).

Bayesian models of perception predict that both such stimuli would be perceived as uniform fields because they would correspond to the most likely cause of the neural activation (Brown and Friston 2012). This proposal is agnostic, however, with respect to whether the underlying neural representations generate retinotopically-distributed activity isomorphic with the fill-in percept at a subsequent stage (Anstis 2010). Such a proposal is compatible with reports that in cortical area V1 the most frequently encountered color sensitive cells are double-opponent that respond best to isolated chromatic contours but also to the edges of uniform color and luminance surfaces (Friedman, Zhou, and von der Heydt 2003, Johnson, Hawken, and Shapley 2001). These cells show response profiles that are stronger for stimulus edges than uniform surfaces (Friedman, Zhou, and von der Heydt 2003), and, in fact, optical imaging experiments using voltage sensitive dyes in macaque (Zweig et al. 2015) and functional imaging in humans (Cornelissen et al. 2006) demonstrate that the response profiles in area V1 for uniform luminance or chromatic surfaces are dominated by edge responses.

In area V1, it is conceivable that surface- and edge-dependent percepts could be processed through distinct neural channels. Double-opponent cells in area V1 could carry information about both isolated chromatic and luminance edge-transients, such as those that generate filling-in, as well as the edges of uniform surfaces while single-opponent cells would respond optimally to the interior region of uniform fields. Single-opponent receptive fields display different spectral sensitivities in spatially distinct excitatory and inhibitory regions, thereby giving rise to spectrally dependent responses to uniform fields. Such an organization would be consistent with theories of visual processing that propose independent surface and edge processing in form perception (Pinna and Grossberg 2005). While the population of single opponent cells in V1 is estimated to be less than half that of the double-opponent cells, this may simply reflect differences in the sampling requirements for detecting uniform surfaces and edges (Schluppeck and Engel 2002). While area V1 has the neural machinery to support both edge-induced and surface-dependent color, here we ask to what extent the mechanisms underlying these perceptual phenomena are dependent on distributed neural activity across cortical areas, and more particularly by distinct inter-areal networks.

The edge-induced, filling-in phenomenon known as the Watercolor Effect (WCE) is ideal for investigating the networks involved in contextual effects (Figure 2a). In the WCE, the color of the inner contour of a pair of distant, chromatic contours modifies the appearance of an interior region that is physically identical to the background in such a manner that it appears as a uniform, desaturated hue (Pinna, Brelstaff, and Spillmann 2001). The WCE propagates over too large a visual angle to be attributed to light spread in the eye. However, to date, investigation of the WCE has provided little evidence concerning the neuronal underpinnings of the phenomena. Importantly, based on theoretical and psychophysical observations (Devinck et al. 2014), the thin chromatic edge-transient that induces the WCE is expected to preferentially activate edge-sensitive double opponent cells in area V1 rather than surface responsive single opponent cells. While Coia et al. (2014) found that the visual evoked potential (VEP) correlates with psychophysical responses for the WCE, insufficient resolution of the VEP make it impossible to localize the areas implicated in the processing. Finally, both the large extent of the filling-in and the sensitivity of the phenomenon to the curvature of the inducing contours (Gerardin et al. 2014) point to the involvement of cortical areas beyond V1 and V2.

**Figure 2:**
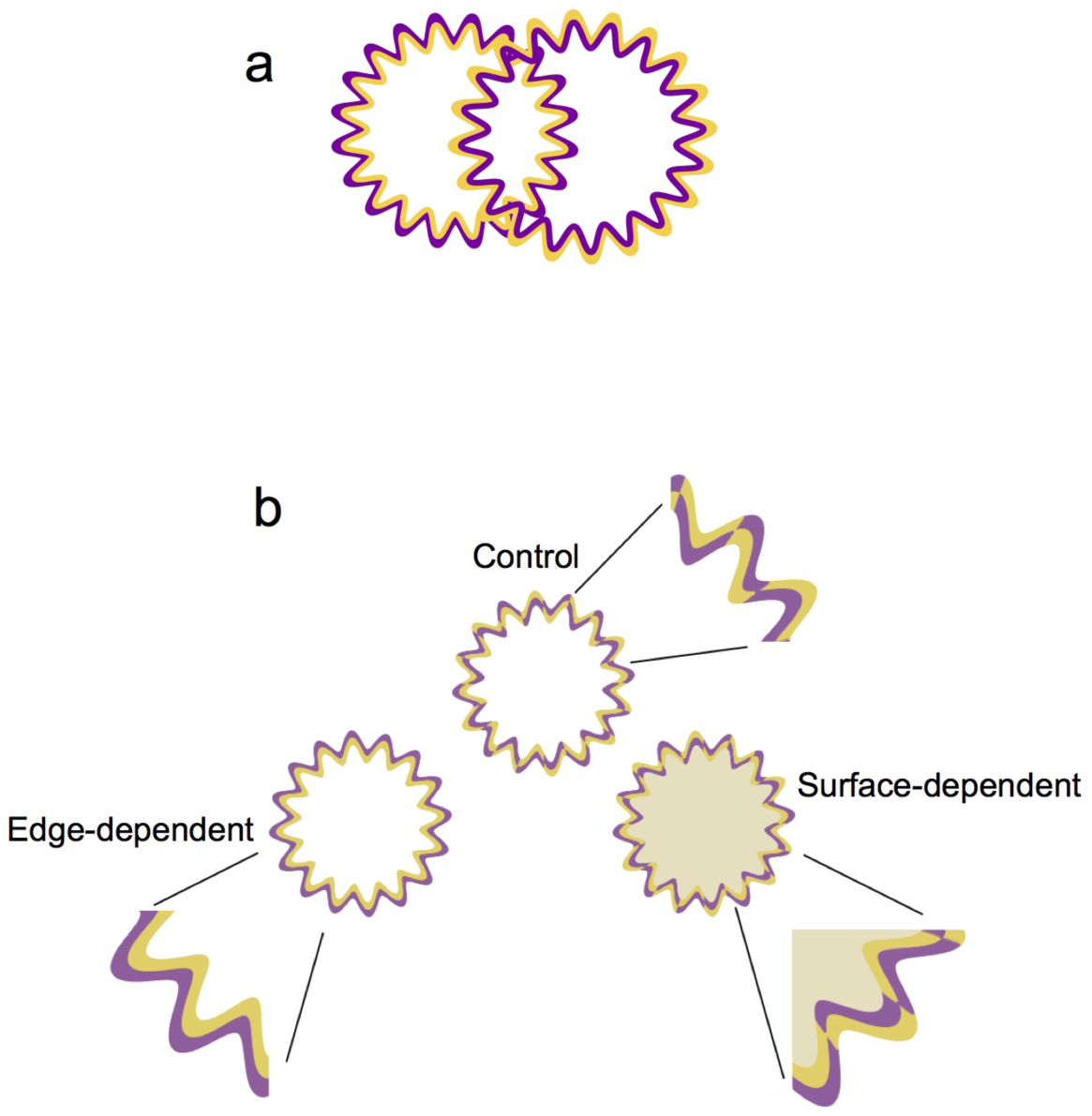
Watercolor effect demonstration and fMRI stimuli used in the main experiment. a. Stimulus demonstrating the WCE. Two intersecting circular chromatic contours, each with its radius frequency modulated, thereby defining three regions. In the left region, the interior region takes on the hue of the bright orange interior contours showing the WCE while in the right, the dark purple contours generate no filling-in. The region of intersection that is enclosed by both types of contours is described by some observers as showing a weaker filling-in color. b. Examples of the three classes of stimuli used in the fMRI experiment. Beside each stimulus an enlarged detail of the contour is drawn to illustrate its structure Left: Edge-dependent stimulus inducing the perception of the WCE, as defined from (Gerardin et al. 2014); Center: The control stimulus is composed of braided contours and demonstrates a decrease in strength of WCE filling-in; Right: The surface-dependent stimulus was used to match the effect of the WCE filling-in, in which the interior chromaticity was defined from each subject’s matches (from a preliminary paired comparison experiment inside the scanner). These examples are schematic in terms of the width and color of the contours and interior region, which are enhanced here for visibility of the stimulus structure used.

Here, we used functional magnetic resonance imaging (fMRI) combined with Multi-Voxel Pattern Analysis (MVPA) in order to localize those cortical areas associated with edge-induced, color filling-in of the WCE phenomenon and those responding to a matched uniform chromaticity. To validate the functional imaging findings, we compared the blood oxygen-level dependent (BOLD) signal with the psychophysically estimated strength of the WCE. The MVPA results demonstrate that neural activity supporting the WCE is distributed across multiple hierarchical levels and streams in the visual system, but only a small set of higher order dorsal areas are related to the perceptual strength of the phenomenon. Combining Dynamic Causal Modeling (DCM) with Bayesian Model Selection, shows that the areas most associated with the WCE exert a feedback modulation of area V1 and the ventral stream, and the directions of these modulations are reversed for the matched uniform chromaticity. The results suggest that the WCE color filling-in depends, at least in part, on a dorsal stream influence on ventral color areas. They support the idea that edge-induced filling-in and surface-dependent color are processed through distinct networks across the cortical hierarchy engaging ascending and descending pathways.

## 2. Material and Methods

### 2.1. Observers

Sixteen observers (10 female, mean±SD age: 28±4 years) participated in the study. All observers had normal color vision (Ishihara test and Panel D15), normal or corrected to normal vision and were all right-handed. The study was approved by the local ethics committee (ID-RCB 2012-A000123-40) and all participants gave written informed consent.

### 2.2. Stimuli

The stimuli were created with Matlab R2010b (Mathworks, MA., U.S.A.), and displayed with the PsychToolbox extensions (Brainard 1997, Pelli 1997). Spectral and luminance calibrations of the display were performed with a PR-650 SpectraScan Colorimeter (Photoresearch) and used for screen gamma-correction in stimulus specification. All stimuli were displayed on a white background (580 cd/m2, CIE xy = 0.29, 0.30). Stimuli in the main experiment were defined by a virtual circle (8 degree diameter, i.e., 4 degree eccentricity when centrally fixated) whose radius was modulated sinusoidally. A subset of observers was also tested independently using a 4 degree diameter stimulus allowing comparison to conditions from previous psychophysical studies (Devinck et al. 2014, Gerardin et al.2014). The 4-degree measures also served as a test for repeatability of the pattern of results. The contour pairs that defined stimuli in all conditions had a combined width of 16 min, i.e., 8 min each. The classical WCE was generated with a pair of continuous adjacent contours and will be referred to as the *test or edge-dependent* condition (Figure 2b, left). The outer contour appeared purple (CIE xy = 0.32, 0.19) and the inner orange (CIE xy = 0.48, 0.34). Control stimuli were defined by interleaving the interior and exterior contours in a braid (Figure 2b, center) (Devinck and Knoblauch 2012). An additional stimulus was tested, referred to as the *surface-dependent* condition, in which the chromaticity of the interior of the control stimulus was modified to match the fill-in color of the edge-dependent condition (Figure 2b, right). The matching procedure is described below. All stimuli were specified in the DKL color space (Derrington, Krauskopf, and Lennie 1984) with purple and orange contours at azimuths of 320 and 45 deg, respectively. Additional information is described in the *Stimuli and conditions* paragraph in the section fMRI Design and procedure: the WCE experiment.

### 2.3. Psychophysical Procedures

Prior to the collection of fMRI data, all observers performed the Maximum Likelihood Difference Scaling (MLDS) task, (Knoblauch and Maloney 2012, 2008, Maloney and Yang 2003) in order to measure the perceived magnitude of the WCE from the stimuli *in situ*, in the scanner, for each observer, following the procedure introduced by Devinck and Knoblauch (2012). To confirm that observers responded according to the fill-in color and not on the basis of other stimulus feature(s), equal numbers of test and control stimuli were interleaved in the session. On each trial, a randomly selected triad of either test or control patterns was presented with three luminance elevations of the orange contour (a, b, c) chosen from a series of 10, with a < b < c. Stimulus b was always the upper stimulus in the middle, and stimuli a and c were randomly positioned below on the left or right side, respectively (Figure 3a). In each session, there was a random presentation of the 10!/(3! 7!) = 120 unique triads from the series of 10 luminances of the orange contour, equally-spaced in elevation from the equiluminant plane in DKL space from 0.1 to 0.9, for both test and control stimuli triads, thus yielding a total of 240 presentations. Frequency and amplitude of the contour were set to 20 cycles per revolution (cpr) and 0.2, respectively, with values shown previously to generate a strong WCE (Gerardin et al. 2014). On presentation of a triad, the observer was instructed to fixate each pattern and to choose which of the two bottom patterns (left or right) was most similar to the upper pattern with respect to the color of its interior region. The observer’s response initiated the next trial. No feedback was provided to observers. Difference scales for test and control stimuli were estimated from the session by maximum likelihood using the *mlds* function from the **MLDS** package (Knoblauch and Maloney 2008) in the open source software R (RCoreTeam 2015). The scales estimated are based on a signal detection model of the decision process and have the property that equal scale differences are perceptually equal, i.e., they are interval scales. When parameterized so that each response has unit variance, the scales can be interpreted in terms of the signal detection parameter *d’* (Knoblauch and Maloney 2012). Additional details on the procedure and modeling can be found elsewhere (Devinck et al. 2014, Devinck and Knoblauch 2012).

**Figure 3:**
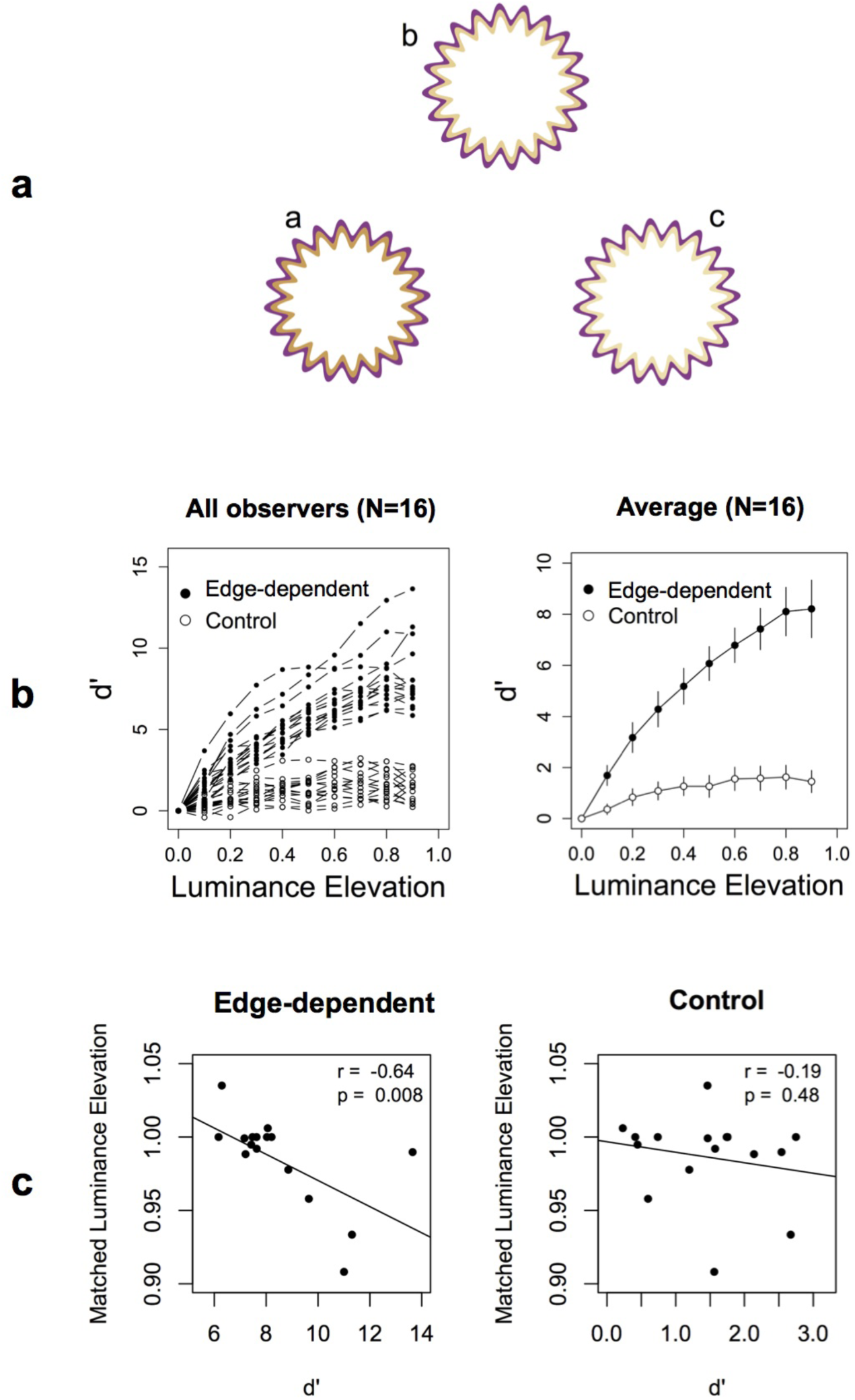
Psychophysics: stimulus configuration and estimated perceptual scales. (a) Example of the triad configuration used in the MLDS task performed in the session before scanning. A triad consisted of either edge-dependent (WCE) or braided control stimuli with three unique ordered luminance elevations (a, b, c, selected randomly from 10 elevations in total) of the interior orange contour. Stimulus b was always the upper stimulus in the middle, and stimuli a and c were randomly positioned on the left or right. The observer had to fixate each pattern until he/she could choose which of the two bottom patterns (left or right) was most similar to the upper pattern with respect to the color of its interior region. (b) Left, all estimated response scales from the MLDS procedure for the sixteen observers as a function of luminance elevation for edge-dependent stimuli (black) and control stimuli (white). The signal detection model used to fit the observer’s data permits the ordinate values to be expressed in terms of the signal detection measure d’. Right, average response scales for the sixteen observers as a function of luminance elevation for edge-dependent and control stimuli with 95% confidence intervals. (c) Correlations of MLDS results with paired-comparison matches. Each point indicates for each observer the luminance elevation that best matched the edge-dependent stimulus (WCE) plotted against the peak MLDS response for edge-dependent (left) and for control stimuli (right).

A paired-comparison experiment was performed to estimate a uniform chromaticity that matched the appearance of the WCE filling-in color. Observers were tested in the scanner prior to scanning. In preliminary observations, we found that the fill-in appearance could be closely matched by simply varying the luminance elevation at constant length of the color vector in DKL space for the interior region. Because the vector length is held constant, the length of the projection on the equiluminant plane decreases with increasing elevation from the plane, thereby reducing colorimetric purity of the stimulus. Observers were presented simultaneously with a test and a control stimulus (braided contour) for which the appearance of the interior of the latter was controlled by adjusting the luminance elevation in DKL color space in the manner described above. Observers judged which interior region of the pair was more orange and the match was assigned to the value at which the choice probability was 50%.

### 2.4. fMRI Design and procedure

Observers were presented sequences of stimuli selected from the three conditions described below. To control for attentional factors, during all functional MRI experiments, observers performed a task requiring them to detect a change in orientation of the fixation cross. Observers were instructed to press a button when they detected a change in the fixation cross from ‘+’ to ‘x’. Performance for this task was 79% ±3% (SD) correct. All stimuli were back-projected using a video-projector on a translucent screen positioned at the rear of the magnet. A subject in the scanner, viewed the screen at a distance of 122cm via a mirror fixed on the head coil.

#### 2.4.1. Stimuli and conditions

Three conditions were used. *a) Edge-dependent stimuli*: Based on our previous studies (Gerardin et al. 2014) and the psychophysical experiments performed in the scanner, stimuli were chosen to generate a strong WCE. Three contour frequencies (16, 18 and 20 cpr) were used, with two contour amplitudes (0.16 and 0.20) and with three luminances of the orange contour (elevations of 0.7, 0.8 and 0.9 in DKL space) (Figure 2b left). *b) Control stimuli:* Control stimuli were identical except that the contours were interlaced or braided and generated little filling-in (Figure 2b center). *c) Surface-dependent stimuli:* These stimuli had braided contours similar to the control, but the region interior to the contours was set to a uniform chromaticity and luminance that matched the interior, fill-in color of the edge-dependent stimuli (Figure 2b right), based on a paired-comparison experiment performed inside the scanner for each subject, described above. These stimuli were designed to generate the same appearance as the uniform interior of the WCE stimulus but with the bounding contour of the control stimulus.

Table 1 shows the pattern of feature differences among the three stimulus conditions that we used and on the basis of which we can make comparisons and inferences about the role of particular Regions of Interest (ROIs) on edge and surface dependent color appearance. Because each comparison varies for two features, no comparison uniquely defines the information treated by a ROI, instead the interpretation of the source of the activation is constrained by the combined classification across all three comparisons.

**Table 1:**
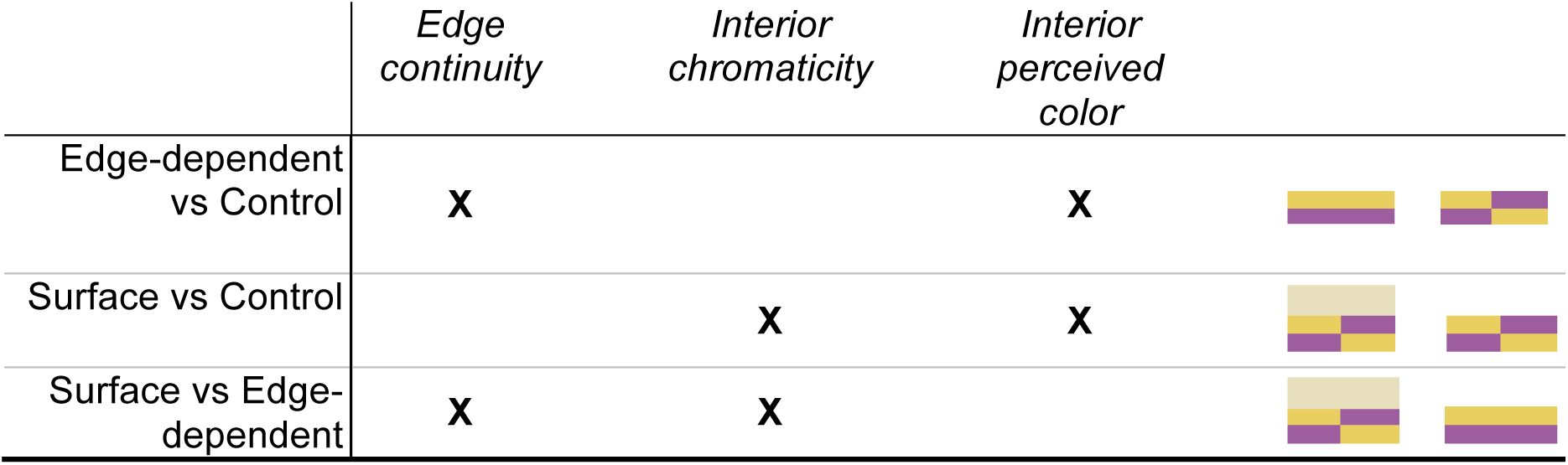
Stimulus and response differences tested by classifiers.

|  | *Edge*<br>*continuity* | *Interior*<br>*chromaticity* | *Interior*<br>*perceived*<br>*color* |
| --- | --- | --- | --- |
| Edge-dependent<br>vs Control | **X** |  | **X** |
| Surface vs Control |  | **X** | **X** |
| Surface vs Edge-dependent | **X** | **X** |  |

The X’s indicate the features on which the comparisons differed and also symbolized by the contour pairs in the last column. The pattern of responses across the three classifiers and not a single classification constrains the specificity of the region tested.

For example, significant differences for the edge-dependent vs. control comparison could result from either the local difference in edge continuity of the inducer or from the difference in perceived color of the interior region. However, if the same ROI only weakly differentiates the surface-dependent vs. edge-dependent, then the influence of the edge continuity on the response can be excluded since these stimuli also differ with respect to that feature. In this case, we infer that the area’s response is implicated in the treatment of filling-in. If in addition the ROI weakly differentiates surface vs. control conditions, then it implies that the ROI’s response does not depend strongly on the surface chromaticity since that feature is common to the two comparisons showing weak classification. On the other hand, if an ROI strongly makes a strong distinction between the edge-dependent vs. surface and the surface vs. control conditions but not the edge-dependent vs. control, then we can conclude that it responds on the basis of a purely stimulus bound feature, the interior chromaticity difference, since that is the only common feature.

#### 2.4.2. Procedure

Each observer was tested in one session comprising six scanning runs. Each run consisted of 12 blocks plus 5 fixation intervals (including one at the start, one at the end, and one between each 3-block sequence) and lasting for 272 sec. Three conditions were tested: a) stimuli inducing WCE (edge-dependent), b) null filling-in stimuli (control) and c) surface-dependent stimuli. Each block comprised 20 stimuli of one condition and was repeated four times. Block presentation was randomized. Each stimulus was presented for 500 ms followed by a blank interval (300 ms).

### 2.5. fMRI Data Acquisition

All experiments were conducted using a 3-Tesla Philips Achieva MRI scanner at the Grenoble MRI facility IRMaGE, France. In each session for each individual, a high-resolution T1-weighted structural image (3D TFE sequence, acquisition matrix 240 × 256 × 180, TR/TE: 25/2.3 ms, flip angle 9°, 1 × 1 × 1mm resolution) and series of T2*-weighted functional images (EPI MS-FFE, acquisition matrix 80 × 80, 30 slices, TR/TE: 2000/30 ms, flip angle 80°, acquisition voxel size 3 × 3 × 2.75mm, reconstructed voxel size 3 × 3 × 3mm) were collected with a 32 channel SENSE head coil.

In order to control for fixation stability, the position of the left eye over the course of all experiments was monitored with an ASL EyeTracker 6000. No systematic deviations from the fixation point were observed and no data were excluded from the analysis.

### 2.6. fMRI Data Analysis

The fMRI data were analyzed using Brain Voyager QX (Brain Innovations, Maastricht, The Netherlands). Preprocessing of the functional data consisted of slice-scan time correction, head movement correction, temporal high-pass filtering (2 cycles) and linear trend removal. Individual functional images were aligned to each corresponding anatomical image that was used for 3D cortex reconstruction, inflation and flattening. The volume time-course datasets were convolved with a canonical hemodynamic response function. No spatial smoothing was applied except for the group whole brain analysis (Gaussian filter; full-width at half maximum: 6 mm kernel) after normalization of all data to a common referential (Talairach space). Functional data analysis was first performed using a general linear model (GLM). Such an analysis was realized on the entire brain and in individually mapped regions of interest. Subsequently, Multi-Voxel Pattern Analyses (MVPA) were performed on each individual with Matlab R2010b (www.MathWorks.com) and SVM^light^ (Joachims 1999). Finally, Dynamic Causal Modelling (DCM) analyses were performed using Matlab (R2014a-with SPM12 (6906) with supplementary analyses using the open source software R (R Core Team, 2017). The three approaches used for fMRI data analysis and how they are related to each other are schematized in Figure 4.

**Figure 4:**
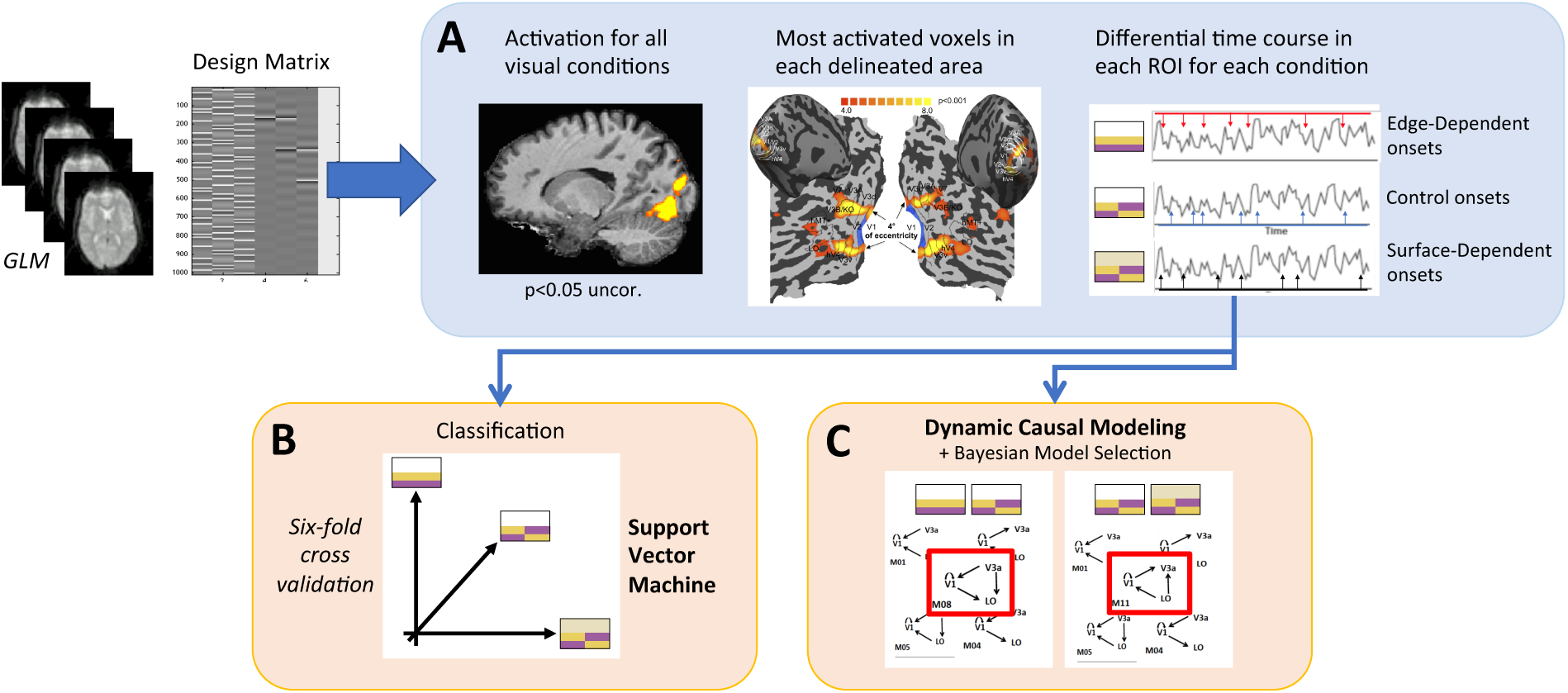
Image processing steps for each subject. **A.** Selection and Time series extraction. After General Linear Model (GLM) estimation, the most activated voxels for all visual conditions were selected in each delineated area and their time course extracted for each condition. Stimulus conditions in right panel are indicated symbolically from top to bottom as continuous contours, braided contours and braided contours with added uniform chromaticity. **B.** Classification. A Support Vector Machine classifier was trained based on the mean z-normed activation of each area in response to each block to discriminate between conditions using a six-fold cross-validation scheme. **C.** DCM analysis. The eigenvectors of selected area in response to each condition were used as input (stimulus comparisons shown above models) for the evaluation of connectivity modulations (12 possible models). These models were then compared within condition with Bayesian Model Selection. Red rectangles indicate the selected model for edge vs control (left) and surface vs control conditions.

#### 2.6.1. General Model Analysis

For each participant, all three conditions (edge-dependent, control and surface-dependent) were modeled as three regressors constructed as boxcar functions convolved with a canonical hemodynamic function. Parameters obtained from movement correction were added to the design matrix as nuisance covariates. Contrast images were computed relative to the fixation condition for all visual conditions and each stimulus condition separately. These contrasts were used to select the most activated voxels in each ROI. Once voxels were selected, the MVPA was run on the raw data (see Figure 4A).

#### 2.6.2. Mapping Regions of Interest

In a second session, for each observer, we identified: 1) retinotopic areas V1 and V2, 2) the human V4 (hV4, also identified by retinotopic mapping) and Lateral Occipital complex (LO) and 3) motion-related areas (V3B/KO, hMT+/V5). We functionally localized early visual areas V1 and V2, dorsal retinotopic areas V3, V3A and V7 and ventral retinotopic areas V3v and hV4 based on standard retinotopic mapping procedures (Warnking et al. 2002, DeYoe et al. 1996, Sereno et al. 1995, Engel et al. 1994). We also functionally localized LO (Kourtzi and Kanwisher 2001), V3B/KO (Dupont et al. 1997), and hMT+/V5 (Tootell et al. 1995) with a block related paradigm detailed below (Figure 5).

**Figure 5:**
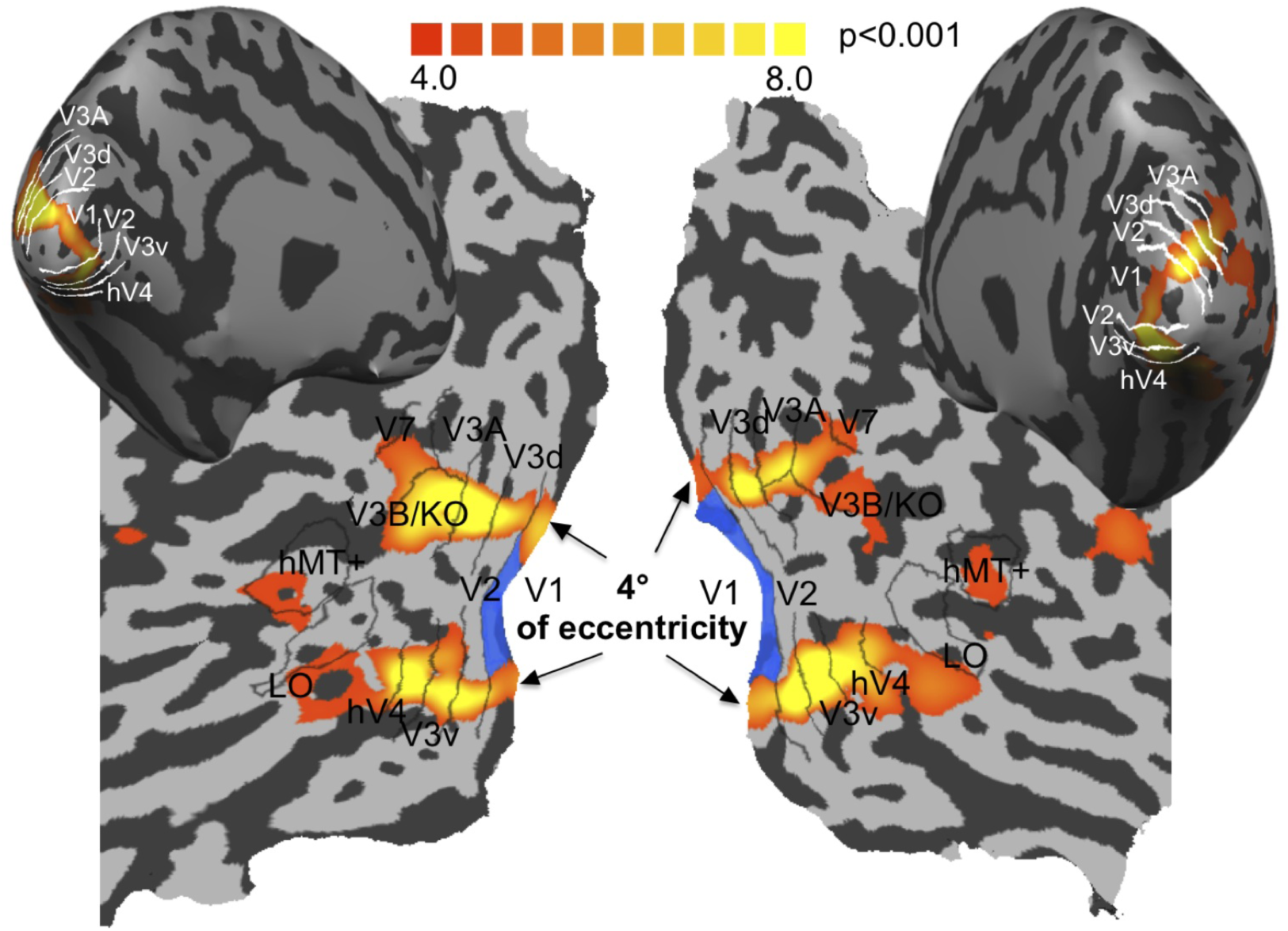
ROIs delineation. Mapping of Regions of Interest (ROIs). We identified: 1) retinotopic areas, 2) the Lateral Occipital complex (LO) and 3) motion-related areas (V3B/KO, hMT+/V5).). Functional activation (from red to yellow, for low to high activation) from the braided control (4 deg eccentricity) versus fixation conditions was projected onto each inflated or flattened hemisphere (Left: left hemisphere; Right: right hemisphere for one subject). The blue shading in area V1 indicates the retinotopic regions interior to the activation by the 4-degree eccentric contours of the control stimulus, based on the eccentricity maps obtained from the retinotopy mapping experiment. This defines the region interior to the stimuli from which the voxels were defined for the analyses in order to exclude voxels responding directly to the contour.

For retinotopic areas, we computed correlation analyses with a sinusoidal function with 16 lags to obtain power and phase maps for both the eccentricity and the polar mapping. Phase maps were thresholded at a correlation of 0.2 and projected on the cortical flat maps. The borders between visual areas were identified as phase reversals on the polar phase maps, with simultaneous visualization of the eccentricity map to ensure that the borders ran perpendicularly to the eccentricity gradient. We identified V1 and V2, dorsal areas up to V7 and ventral areas up to hV4. In addition, for area V1, a set of voxels was selected that was retinotopically interior to those significantly activated by the contours of the control stimulus (4-degree eccentricity, blue shading in Figure 5) for use in the subsequent MVPA analysis. This selection was based on the data from the retinotopic mapping experiments, i.e., eccentricity maps defined using expanding concentric rings. This allows excluding pattern classification based on voxel response to the contour. Subsequently, additional control analyses were performed to evaluate the role of the contour in classification for the extrastriate visual areas.

Using the general linear model (GLM) including fixation periods and movement correction parameters as covariates of no interest, we identified LOC, hMT+/V5 and V3B/KO. LOC was defined as the set of contiguous voxels in the ventral occipito-temporal cortex that showed significantly stronger activation (t(165)>4.0, p<0.001 uncorrected) to intact compared to scrambled images of objects (Kourtzi and Kanwisher 2001). hMT+/V5 (Tootell et al. 1995) was defined as the set of contiguous voxels in the lateral occipito-temporal cortex that showed significantly stronger activation (t(165)>4.0, p<0.001 uncorrected) to moving compared to static random dots. V3B/KO was defined as the set of contiguous voxels anterior to V3A that showed significantly stronger activation (t(165)>4.0, p<0.001 uncorrected) to random-dot displays that defined relative rather than transparent motion (Dupont et al. 1997).

#### 2.6.3. Multi-voxel pattern analysis

Multi-voxel Pattern Analysis (Haynes and Rees 2005, Kamitani and Tong 2005), using linear Support Vector Machine (SVM) classifiers followed by cross-validation procedures, was used with Matlab R2010b (www.MathWorks.com) (Gerardin, Kourtzi, and Mamassian 2010) and SVM^light^ (Joachims 1999). For each Region of Interest (ROI: retinotopic areas, LO, V3B/KO and hMT+/V5), voxels were sorted according to their response (t-statistic maps based on the GLM) to all stimulus conditions compared to fixation baseline across all experimental runs. The number of voxels was selected across individual ROIs and observers by restricting the pattern size to those voxels that showed a significant (p<0.05 uncorrected) t value. One hundred voxels from both hemispheres were included for each ROI and subject. For area V1, voxels were chosen to be at retinotopic eccentricities inferior to those activated directly by the control stimulus contour at 4 degrees (blue region of V1 in Figure 5). For other visual areas, it was not possible to exclude such voxels for all observers and maintain 100 voxels per ROI. As an additional test of a possible influence of the contour on the activations, we performed a second analysis in which the number of voxels was reduced per area on an individual basis to the maximum number excluding the activation by the control contour. To account for the differences in voxel numbers between areas and observers, we report weighted means and confidence intervals based on weighted standard deviations for these results with the voxel numbers used as weights. Each voxel time course was normalized (z-score) separately for each experimental run in order to minimize baseline differences. For each subject and each condition, all time series data points of the corresponding experimental block were averaged in order to generate the data vectors for the multivariate analysis. Data vectors were selected according to the comparison of interest and split into a training sample comprising the data of five runs and a test sample comprising the remaining run. A six-fold cross-validation was performed leaving one run out (test sample). For each subject, accuracy rates were averaged (number of correctly assigned test patterns/total number of assignments) across cross-validation runs. Statistical significance across conditions was evaluated using linear mixed-effects models with observer treated as a random effect (Pinheiro and Bates 2000). A recent study questioned the use of parametric significance tests with MVPA based on simulations that revealed skewness of the null-distribution (Jamalabadi et al. 2016). Diagnostic plots of the residuals from the mixed-effect model fits (residuals vs fitted values, and quantile-quantile plots, S1), however, revealed no systematic deviations from model distribution or variance assumptions that would warrant discounting the model or preferring transformed values of the response variable.

#### 2.6.4. Control analyses for the MVPA

Control analyses were performed with permutation tests to evaluate whether the observed classification accuracies observed were due to chance (Etzel 2017, Etzel and Braver 2013). The MVPA analyses were run with randomly assigned category labels to each activation pattern for 1000 repetitions per subject. For each permutation, the classification accuracies were averaged across subjects. The probability of observing a value equal or higher than the average observed value was calculated from the distribution of permuted averages as the achieved significance level of the test (Efron and Tibshirani 1994). Ninety-five percent confidence limits for the mean predictions under the permutations were estimated from the 2.5 and 97.5 quantiles of the permutation distributions.

#### 2.6.5. Dynamic Causal Modeling for BOLD responses

Using Dynamic Causal Modeling (DCM), we explored changes in the effective connectivity, i.e. the inferred influence exerted by one region on the others, and how information is propagated through these regions (see (Friston 2011) for details about the operational distinction between functional and effective connectivity) in response to our protocol. DCM considers the brain as a deterministic system whose response in a region or part of a cortical network, is determined by activity in other regions. We defined causal models of connectivity between selected ROIs to make inferences about underlying mechanisms expressed in our BOLD data. These models included 1) intrinsic (or endogenous) connections that quantify the effective fixed connectivity between the model nodes, i.e. the changes in activity in a target region when activity in the source region changes (matrix A in DCM), 2) stimulus-related inputs that define how the model responds to task-related inputs (matrix C in DCM), and 3) possible input modulations that define how the effective connectivity is influenced by experimental factors (matrix B in DCM). Following model estimations, DCM provides a Bayesian method (Bayesian Model Selection, BMS) for selecting the best model of coupling that explains connectivity changes underlying task-related brain responses (Friston, Harrison, and Penny 2003). Note that effective connectivity between regions does not imply underlying direct anatomical connections (Friston 2011).

##### 2.6.5.1. Model structure

Following BMS, each model is attributed a probability according to its power to describe the data. These probabilities sum to one, so the number of models in the model space must be limited to prevent excessive dilution of the maximum probability. One way to achieve this is to limit the number of areas considered. In our case, three areas were considered in each model definition because it is the lowest number necessary to translate our hypothesis concerning the connectivity modulations between dorsal and ventral streams. For example, region V1 (as receiving visual input), region V3A (that best classified edge-dependent stimuli) and region LO (that best classified surface-dependent stimuli). As early visual areas are strongly interconnected, we considered all possible connections between and within areas in our endogenous matrix. Our control stimuli entered all models as a driving input to V1.

##### 2.6.5.2. Connectivity Modulation

We designed a model space in terms of which a subset of connections was modulated by the edge-dependent or the surface-dependent stimuli. We reasoned that both types of stimuli have a differential effect on endogenous connections of the network for each subject. We did not consider bidirectional modulations because we were only interested in the strongest direction. V1 was always connected to the two other areas in all models, but we included the hypothesis that there was no differential modulation between them. Because of the importance of V1 intrinsic horizontal connections for contour integration (Stettler et al. 2002), for each model we included a self-modulation of V1. This resulted in the model space described in Figure 6a and Supplementary Figure S2a with 12 possible modulations.

**Figure 6:**
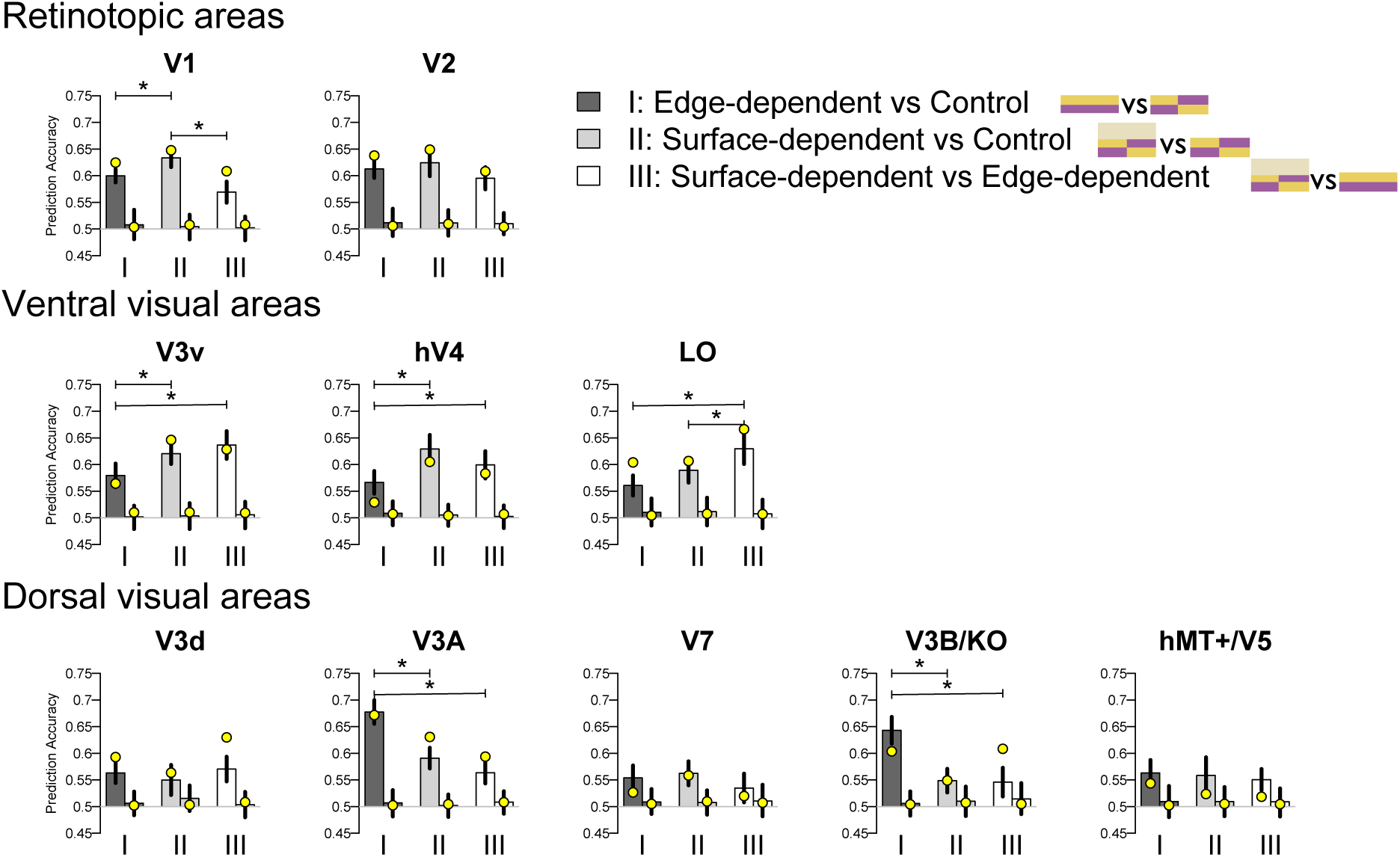
Multi-Voxel Pattern Analysis (MVPA) from fMRI data. Classification accuracies across the 10 regions of interest for the three classifiers: (I) Edge-dependent vs. control (dark grey); (II) Surface-dependent vs. control (grey); (III) Surface-dependent vs. edge-dependent (white). Mean classification accuracy is based on 100 voxels per area in both hemispheres. Each bar plot shows results for 3 pairs, means of observed values across observers (left) and means based on permutation tests (n = 1000) (right), of classifiers. Error bars indicate 95% confidence intervals. Significant differences between the classifiers (p < 0.05) controlled for multiple comparisons (Tukey test) are indicated by horizontal bars and star. The yellow points indicate average results of a repeated experiment on three of the observers using a 4 deg stimulus.

##### 2.6.5.3. Model comparison

All twelve models were applied successively with the functional datasets from each subject and each condition (edge-induced or surface-dependent). They entered a Bayesian model selection procedure that identified which of the competing models best predicts each dataset. For each model, the model evidence, i.e. the probability of observing the measured data given a specific model, was computed based on the free energy approximation. Model evidence was used by the Bayesian model selection procedure to rank the models. The protected exceedance probability defined the relative superiority of a model, given the group data (Rigoux et al. 2014). For group-level Bayesian model selection we then considered a Random-effects (RFX) analysis to account for between-subject variability (Stephan et al. 2009).

## 3. Results

### 3.1. Estimated perceived strength of WCE

Perceptual scales were estimated by Maximum Likelihood Difference Scaling (MLDS), method of triads (Knoblauch and Maloney 2008, Maloney and Yang 2003), a powerful method for identification of perceptual correlates of BOLD responses (Bellot et al. 2016, Yang, Szeverenyi, and Ts’o 2008). A sample trial illustrating the stimulus configuration is shown in Figure 3a. The approach is based on the idea that when the perceived filling-in of the upper stimulus is judged equally often to be as similar to the lower left as to the lower right stimulus, then the perceived differences between a and b and between b and c are equal. This allows scale values to be estimated by maximum likelihood based on a signal detection model of the observer decision rule (Knoblauch and Maloney 2012, Maloney and Yang 2003). Individual (left panel) and average (right panel) perceptual scales estimated by MLDS (Figure 3b) display the dependence of the strength of the filling-in on the luminance of the orange contour for test (filled circles) and control (open circles) stimuli. With test stimuli the filling-in response increases with the elevation of the luminance of the interior contour with respect to an equiluminant plane in DKL space, while for the control stimuli responses are attenuated or absent (Devinck et al. 2014, Devinck and Knoblauch 2012). The differences in response magnitude between edge-dependent and braided control stimuli confirms that observers respond to the fill-in color and not the contours, and reveals significant variation in the strength of the WCE across observers. Figure 3c shows for all observers two scatterplots of the luminance elevations that best match the WCE against the peak MLDS responses for the edge-dependent stimulus (left panel) and the control stimulus (right panel). As the luminance elevation of the matched stimulus decreases in DKL space, colorimetric purity increases. Thus, lower luminance matches correspond to a higher matched colorimetric purity and therefore, a stronger perceived fill-in color in the edge-dependent stimulus. The strength of the WCE is significantly correlated with the matched color value (r=-0.62, p=0.01) for the edge-dependent but not for the control stimulus (r = -0.19, p = 0.48). Thus, individual differences in the strength of the WCE covary similarly for both the scaling and matching data.

### 3.2. BOLD activation related to the WCE

A whole-brain analysis using a General Linear Model (GLM) was undertaken so as to detect significant BOLD activation related to the WCE. Figure S2 in Supplementary Material shows the activation in each ROI with respect to the mean response for the fixation condition and normalized by the pooled standard deviation, for each condition. For all areas, ninety-five percent confidence intervals overlap in all conditions, except for LO, V3d and V3A, which show higher activation for the Edge condition that generates a WCE. To detect differential activation, stimulus blocks from each of the three conditions were contrasted with the other two in a group analysis. No significant activation survived when correction for multiple comparisons was applied (p>0.05, Bonferroni corrected for all brain voxels).

### 3.3. Discriminating edge-dependent and surface-dependent processing with MVPA

To obtain a more fine-grained analysis of the contrasts, MVPA was applied to ROIs identified by functional localizers. We assumed that if a ROI is involved in edge- or surface-dependent color appearance, contrasts in the stimuli would generate a significantly higher accuracy than chance according to the logic expressed in the methods with respect to Table 1. Signal-to-noise ratios tested by a linear mixed-effects model across cortical areas and with a random observer intercept showed no significant differences that could bias such comparisons (Likelihood ratio test: 𝒳^2^ 9 = 7.4; *p* = 0.6, Figure S3).

MVPA was used to test whether activity patterns of voxels in retinotopic areas (V1, V2), dorsal areas (V3d, V3A, V7, V3B/KO, hMT+/V5) and ventral areas (V3v, hV4, LO), decode the WCE. As demonstrated previously (Devinck et al. 2014, Gerardin et al. 2014) and confirmed in the psychophysical experiments above, WCE filling-in is absent or strongly attenuated when the stimulus contour is braided. Classifiers were run to identify regions that decode preferentially these particular properties of the edge-dependent stimuli (WCE). Analyses were also conducted to reveal visual regions that classify the surface-dependent stimuli matched in color appearance to the WCE. For each area, three classifiers tested specific hypotheses: (I) that accuracy was higher for the classification of edge-dependent compared to control stimuli; (II) that accuracy was higher for the classification of surface-dependent compared to control stimuli and (III) that accuracy was higher for the classification of edge-compared to surface-dependent color stimuli. All results reported below were corrected for multiple testing (Holm 1979).

Figure 6 shows the classification accuracies (left bar of each of the three pairs in the bar graph for each area with 95% confidence intervals and Supplementary Table S1) with comparisons to the mean value obtained from a permutation test (n = 1000, right bar in each pair with 95% confidence intervals of the permutation distributions) for each of the comparisons among the three stimulus conditions and each ROI. The confidence intervals for all means based on the permutation distributions include the chance level of 0.5. Supplementary Table S2 shows the significance levels after Bonferroni correction of the permutation analyses of the observed classification accuracies. Seven values do not differ significantly from the mean permutation value (Edge-dependent vs Control: hMT+/V5; Surface-dependent vs Control: V3d, V3B/KO, hMT+/V5; Surface-dependent vs Edge-dependent: V7, V3B/KO, hMT+/V5). All other values fell outside the distribution of 1000 permutations. The results suggest that processing of the stimuli tested is distributed across multiple areas. However, not all areas displayed differential classification for the three comparisons. For 4 areas (V2, V3d, V7 and hMT+/V5), the classification accuracies showed no selectivity for the conditions (Table S6, linear mixed effects model: V2: F(2, 30) = 2.96, p = 0.07; V3d: F(2, 30) = 0.87 p = 0.42; V7: F(2, 30) = 1.53, p = 0.23; hMT+/V5: F(2, 30) = 0.28, p = 0.76), and except for area V2, which classified robustly for all three MVPA comparisons, displayed relatively low or non-significant prediction accuracies. Significant differences between the MVPA comparisons were observed in V1 (Table S6, linear mixed-effects model: F(2,30)=16.58, p<0.001) and in ventral stream areas (V3v: F(2,30)=7.61, p<0.01); hV4: F(2,30)=11.78, p<0.001; LO: F(2,30)=17.14, p<0.0001). In dorsal areas, differences between the MVPA comparisons only attained significance for areas V3A (F(2,30)=47.05, p<0.00001) and V3B/KO (F(2,30)=27.27, p<0.00001) (Supplementary Table S6). The yellow points indicate independent replications of the conditions in 3 observers using a 4 degree stimulus. In all cases, they yield the same pattern of results as for the larger stimuli and support the robustness of the findings.

Ventral and dorsal stream areas systematically differ in the profile of classification accuracy across the three MVPA comparisons. Ventral areas (V3v, hV4 and LO) displayed high accuracies for the surface-dependent vs. edge-dependent classifier that additionally were significantly greater than for the edge-dependent vs. control comparison (Table 2, I vs III: Tukey test, V3v: t(885) = -4.3, p < 0.001; hV4: t(885) = -2.5, p < 0.05; LO: t(885) = -5.2, p < 0.001). If the classifiers were driven by the edge continuity, then both of these comparisons would be expected to indicate high prediction accuracy. Areas V3v and hV4 also showed a significantly higher prediction accuracy for the surface-dependent vs. control comparison than for the edge-dependent vs. control (Table 2, Tukey test, I vs II: V3v: t(885) = -3.1, p < 0.01 and hV4: t(885) = -4.7, p < 0.001). This result also argues against the edge continuity as the decoding feature because the edge-continuity is matched for the comparison with higher prediction accuracy (surface-dependent vs control). Instead, the two comparisons differ in the interior chromaticity but not the perceived interior color, suggesting that decoding is based on the former feature. LO shows a similar prediction accuracy profile except that the difference between comparisons I and II does not reach significance (edge-dependent vs control compared with surface-dependent vs control: Table 2, Tukey test, I vs II; LO: t(885) = -2.1, p = 0.08). These findings suggest that the classifications do not depend on either edge continuity or interior chromaticity, since both of these classifiers differ on these attributes and if one were driving the decoding results, then the classification accuracies would differ, too. The highest prediction accuracy occurs, however, for the classification that compares stimuli in which the interior perceived colors match. This would suggest that LO classification depends on the perceived color evoked by the chromaticity difference but not that arising specifically from filling-in.

**Table 2:**
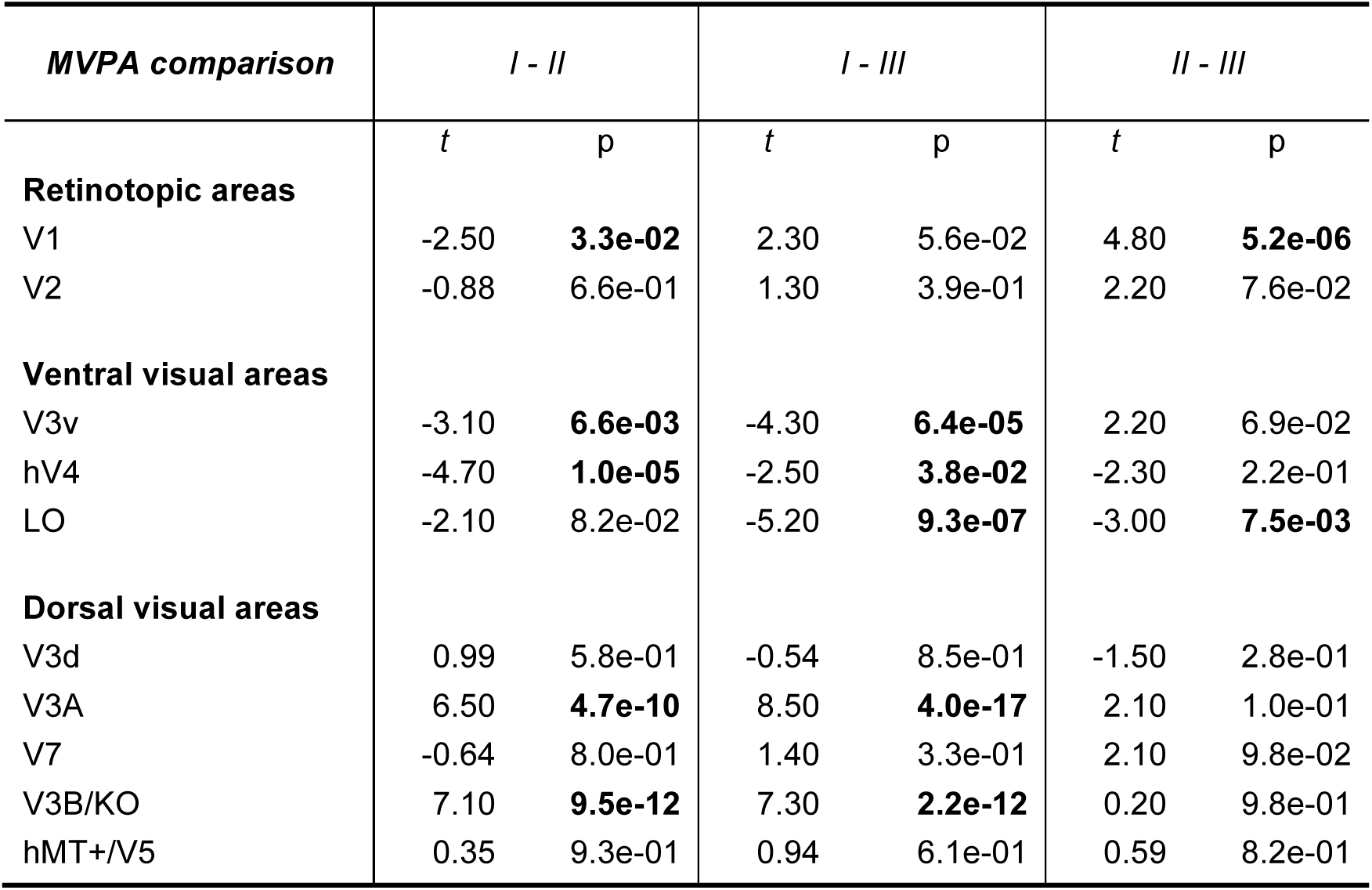
Post-hoc t-tests (Tukey Honest Significant Differences test) for differences of classification accuracy between classification comparisons within areas. *Roman numerals refer to classification comparisons:* (I) Edge-dependent vs. control; (II) Surface-dependent vs. control; (III) Surface-dependent vs. edge-dependent. Each column indicates the t- and p-values for the difference of a pair of classification comparisons. Significance levels for p < 0.05 are indicated in bold.

| ***MVPA comparison*** | ***I - II*** |  | ***I - III*** |  | ***II - III*** |  |
| --- | --- | --- | --- | --- | --- | --- |
|  | ***t*** | ***p*** | ***t*** | ***p*** | ***t*** | ***p*** |
| **Retinotopic areas** |  |  |  |  |  |  |
| V1 | -2.50 | **3.3e-02** | 2.30 | 5.6e-02 | 4.80 | **5.2e-06** |
| V2 | -0.88 | 6.6e-01 | 1.30 | 3.9e-01 | 2.20 | 7.6e-02 |
| **Ventral visual areas** |  |  |  |  |  |  |
| V3v | -3.10 | **6.6e-03** | -4.30 | **6.4e-05** | 2.20 | 6.9e-02 |
| hV4 | -4.70 | **1.0e-05** | -2.50 | **3.8e-02** | -2.30 | 2.2e-01 |
| LO | -2.10 | 8.2e-02 | -5.20 | **9.3e-07** | -3.00 | **7.5e-03** |
| **Dorsal visual areas** |  |  |  |  |  |  |
| V3d | 0.99 | 5.8e-01 | -0.54 | 8.5e-01 | -1.50 | 2.8e-01 |
| V3A | 6.50 | **4.7e-10** | 8.50 | **4.0e-17** | 2.10 | 1.0e-01 |
| V7 | -0.64 | 8.0e-01 | 1.40 | 3.3e-01 | 2.10 | 9.8e-02 |
| V3B/KO | 7.10 | **9.5e-12** | 7.30 | **2.2e-12** | 0.20 | 9.8e-01 |
| hMT+/V5 | 0.35 | 9.3e-01 | 0.94 | 6.1e-01 | 0.59 | 8.2e-01 |

Dorsal stream areas (V3A and V3B/KO) showed a complementary pattern of results in which classification accuracy was highest for the edge-dependent vs. control condition and significantly higher than for the other two comparisons (Table 2, Tukey test, V3A, I vs. II: t(885) = 6.50; p < 0.001; I vs. III; t(885) = 8.50, p < 0.001; V3B/KO: I vs. II: t(885) = 7.10, p < 0.001; I vs. III: t(885) = 7.30; p < 0.001). The high accuracy in these areas for the edge-dependent vs. control comparison supports either a classification on the basis of the filling-in or the edge continuity. The surface-dependent vs. edge-dependent comparison, however, also differed in edge continuity so that the significantly lower response in this case argues in favor of filling-in and against a role for edge continuity. In addition, the significantly lower response of the surface-dependent vs. control comparison argues against a critical role for surface chromaticity *per se* in the classification behavior of these areas.

Areas V1 and V2 showed a similar prediction accuracies for the three classifiers, with higher response for the first two, edge-dependent vs. control and surface-dependent vs. control, but only in area V1 do the differences between comparisons attain significance. This pattern is consistent with processing of the fill-in color and the surface color since the responses are equally strong for the edge- or surface-dependent color and weakest when this attribute is matched. However, the strong activity for all three classifiers could indicate the presence of subpopulations that respond differentially to all of the dimensions along which these stimuli differ.

In order to exploit 100 voxels per area in all observers, the ROIs used for the MVPA analyses other than for area V1 contained voxels that included activation by the stimulus contour. Could the decoding results be due to the contour differences, *per se*. At least two of the comparisons in Table 1 involve pairs of stimuli that differ in contour continuity (I: Edge-dependent vs Control and III: Surface-dependent vs Edge-dependent). Hence, if the contour continuity were the basis of the classification accuracies, we would expect that the classification accuracies obtained for these two comparisons would be correlated. We calculated the Pearson correlations between the prediction accuracies for these two comparisons. Only the correlation for area LO attains significance, and this p-value would not survive a correction for multiple tests (Supplementary Table S3). These findings argue against the contours making a significant contribution to the decoding. We further verified the absence of a role of the contours by determining new ROIs in each area as the maximum number of voxels excluding the contour activation and repeating the analyses. This resulted in unequal numbers of voxels for each area and observer as indicated in Supplementary Table S4. To account for the differences in number of voxels in the analyses, we computed weighted means and standard deviations with the number of voxels as weights. The prediction accuracies for this analysis and the comparisons with the permutation distributions are shown in Supplementary Figure S4 in the same format as Figure 6. The prediction accuracies tend to be a little smaller and the confidence intervals larger as would be expected from an analysis with fewer samples. Several of the lower prediction accuracies now do not differ from chance. However, qualitatively, most of the results show the same trends. The V1 results are the same as those from the previous figure, because the voxels from that area were already chosen to exclude the contour activation zones. Contour exclusion led to the following differences. In area V2, the surface-dependent vs control prediction accuracy is not significant, suggesting that the decoding when the contour is excluded is driven by the contour. In area hV4, only the surface-dependent vs control comparison is significant, suggesting that the decoding is driven by the interior chromaticity. Finally, in area V3v appears to be driven by a combination of the interior chromaticity and the edge continuity. Importantly, the decoding profiles for areas V3A and LO on which subsequent analyses are based are similar for both voxel selections.

### 3.4. Relationship between behavior and neural processing of the WCE

The MVPA analysis reveals candidate areas implicated in the processing of the WCE filling-in, but does not necessarily indicate any relation to perception. To explore such a link, the Pearson product-moment correlations were calculated between the individual classification accuracies for the edge-dependent vs. control MVPA comparison and the peak values for the estimated perceptual scales (d’ from the MLDS task). None of the other comparisons should be related to the perceptual strength of the WCE and with one exception (area hMT+/V5, surface-dependent vs. edge: r = 0.52, p = 0.04), no significant positive correlations were observed. The statistical significance of this case is surprising, given that the prediction accuracy is low (0.55). Figure 7 shows the scatterplots of classification accuracy vs. d’ with regression lines for all ROIs with the correlations and significance levels under the hypothesis of no correlation indicated in each figure. This shows that only areas V3A and V3B/KO showed a significant positive correlation (V3A: r = 0.66, p = 0.006, 95% CI (0.237, 0.869); V3B/KO: r = 0.5, p = 0.049, 95% CI (0.004, 0.794)).

**Figure 7:**
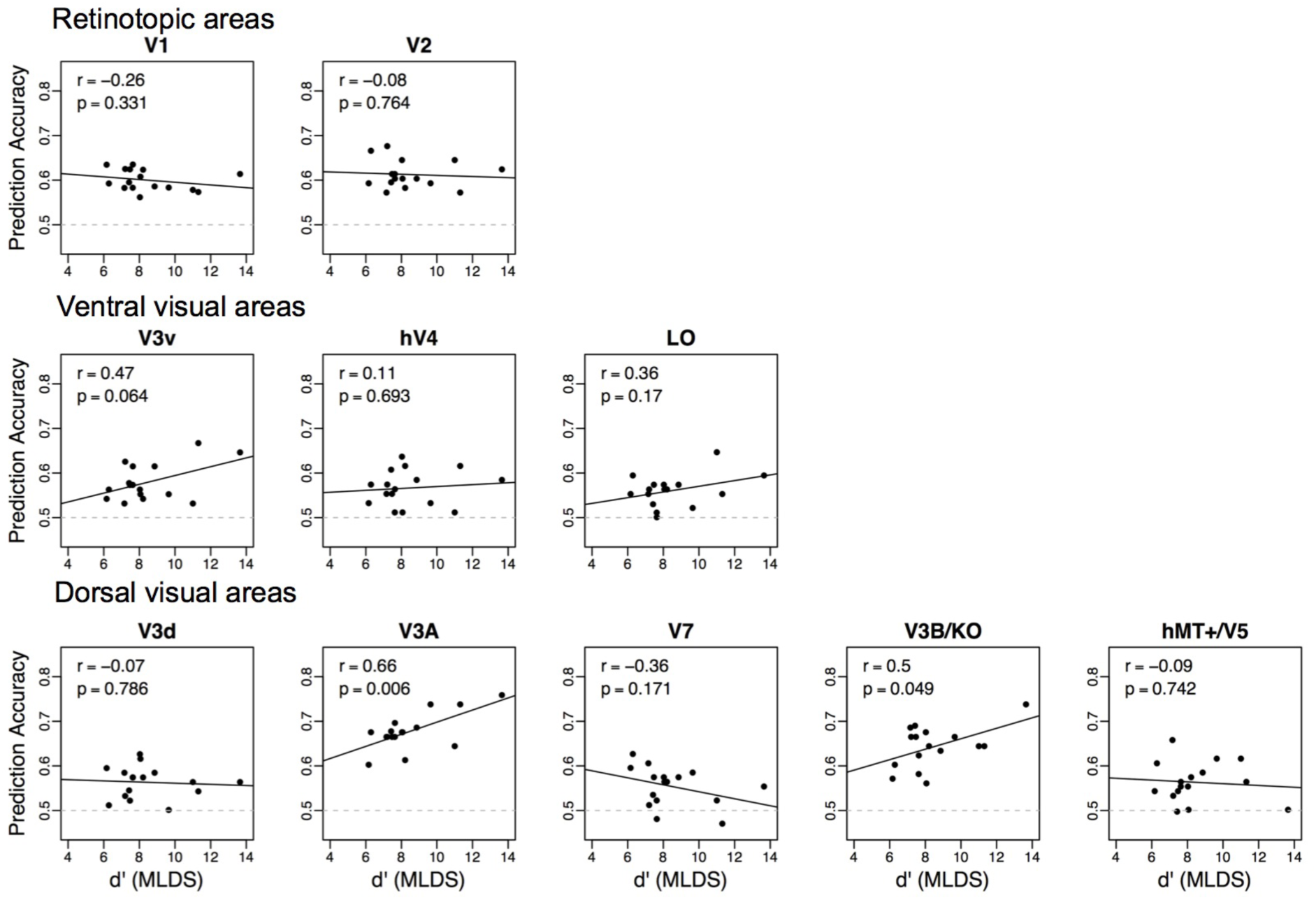
Correlating pattern classification and appearance. Scatter plots of classification accuracies and peak MLDS scale values for all subjects (N=16) for the comparison edge-dependent (WCE) vs. control stimulus for all ROIs with the best-fitting linear regression (solid line). Pearson product-moment correlations and p-values for the hypothesis that the correlation is 0 are indicated within each graph.

The correlation for area V3v is near the criterion of significance used throughout (V3v: r = 0.47, p = 0.064, 95% CI (−0.030, 0.785), and one can legitimately question whether its value differs significantly from that observed for area V3B/KO (Nieuwenhuis, Forstmann, and Wagenmakers 2011). An analysis of covariance to predict classification accuracy with respect to the covariate d’ and the factor Area indicated a significant interaction (F(9, 140) = 2.21, p = 0.025), demonstrating the presence of at least one slope that differed significantly from 0. Inspection of the estimated coefficients for each area revealed significant slopes for areas V3A (t(140) = 2.81, p = 0.006), V3B/KO (t(140) = 2.43, p = 0.02) and V3v (t(140) = 2.04, p = 0.04). Post-hoc tests, however, showed that the slopes of both areas V3A and V3B/KO differed significantly from that of V3v (Tukey Honest Significant Differences test: V3v - V3A: t(140) = -7.4, p < 0.0001; V3v - V3B/KO: t(140) = -4.81, p = 0.0002) while the slope difference between areas V3A and V3B/KO is at chance level (V3A – V3B/KO: t(140) = 2.59, p = 0.2305).

The results of the correlations recomputed with the prediction accuracies based on the contour excluded ROIs were unchanged and significant only for areas V3A and V3B/KO (V3A: r = 0.59, p = 0.016; V3B/KO: r = 0.513, p = 0.042).

### 3.5. Effective connectivity for edge-induced and surface-dependent colors

The above results suggest a role for dorsal areas, particularly area V3A, in the WCE color filling-in. This is surprising because this area, and more generally, dorsal stream areas are little implicated in color processing. We hypothesized that such a role could be mediated through the action of these dorsal stream areas on the ventral stream, which has more typically been implicated in color processing. To evaluate this hypothesis, we used Dynamic Causal Modeling (DCM, (Stephan et al. 2010, Friston, Harrison, and Penny 2003)) and Bayesian Model Selection (BMS) to test the context dependent modulation among areas that showed differential responses to WCE and the uniform added chromaticity. To increase the power of the analysis, we limited evaluation to a triple of areas that included area V1, which represents the visual input at the cortical level, and two areas, V3A and LO, which based on the MVPA analyses, displayed the strongest contrast in their classification profiles with respect to edge-dependent and surface-dependent stimuli. We hypothesized that these areas would be the most likely set of areas to differentiate between the two conditions. We considered models that contrasted the directional modulation of effective connectivity between each pair of the triple when compared with the braided control stimulus. We only tested models with unidirectional modulation of connections between each pair of areas to further reduce the number of models and because we were only interested in the strongest modulation of effective connectivity between area pairs. We considered a space of 12 models (4 patterns of modulation of connectivity of area V1 to V3A and LO times 3 possible relations between V3A and LO), representing plausible effective connectivities between these two areas and V1 (Figure 8a). V1 was always connected to the two areas, but we also included the hypothesis that there was no modulation between V3A and LO.

**Figure 8.**
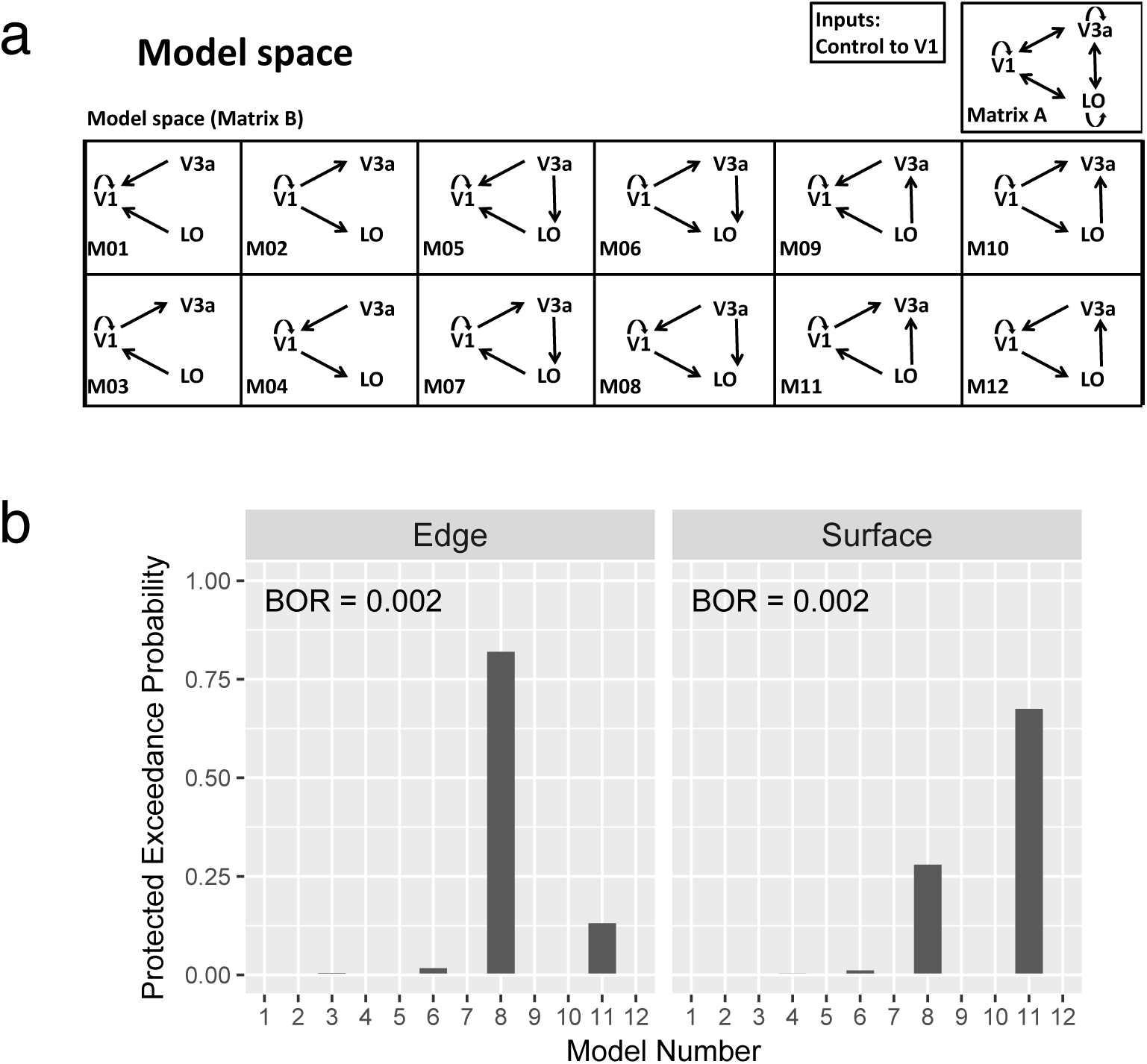
DCM Model space and protected exceedance probabilities for areas V1-V3A-LO. a) Twelve models (M0-M12) of the modulations of effective connectivities between areas V1, V3A and LO were evaluated in the DCM analysis. The intrinsic connectivity (Matrix A) was taken to be complete within and between areas. Activations in response to the Control condition (braided stimuli, no perceived color) were used as input in V1, while activations in response to the Edge dependent or Surface dependent conditions could modulate effective connectivity (Matrix B) according to each model. Each model included self-modulation in V1 and unilateral modulations between V1 and area V3A and between V1 and area LO, yielding 4 possible combinations. Additionally there could either be no modulation between area V3A and area LO (M01 to M04), unilateral modulation from area V3A to area LO (M05 to M08) or unilateral modulation from area LO to area V3A (M09 to M12). b) Protected exceedance probabilities (PEP) for each of the 12 models of modulation of the effective connectivities between the triple of areas indicated in (a) for the edge dependent (left) and surface dependent (right) stimuli. The Bayesian Omnibus Risk (BOR) is indicated in the upper left corner of each plot.

The protected exceedance probabilities (PEP) for the 12 models for each of the two conditions are shown in the bar graphs of Figure 8b for the edge-(left panel) and surface-dependent (right panel) conditions. The Bayesian Omnibus Risk factors (BOR) for the PEP profiles rejected the hypothesis that the models are equiprobable, thus, indicating that at least one differed significantly from the others (BOR = 0.002 for both cases). For the number of observers in our study, PEP values above 0.5 (the disambiguation threshold) are considered strong evidence in favor of the model (Rigoux et al. 2014). Thus, the edge-dependent condition strongly supported model M08 (PEP = 0.82) in which feedback connections from area V3A influence areas V1 and LO, and a feedforward connection from V1 modulates LO. The surface-dependent condition, however, strongly supported model M11 (PEP = 0.68), which displays a complementary set of modulations, i.e., LO exerting a feedback modulation on V1 and V3A, and V1 a feedforward modulation of area V3A.

In the analyses above, areas V3A and LO were selected based on the clear contrast in their stimulus classification profiles. Concomitantly, we should expect that pairs of areas for which the classification accuracy was low and for which the profiles did not distinguish among the stimuli would not show evidence supporting different models of effective connectivity for the edge- and surface-dependent conditions. As a control, we examined a model space for the triple V1-V3d-V7 (Supplementary Figure S5a). Both V3d and V7 showed low classification accuracies (p < 0.6) and displayed no significant differences among the three comparisons (Figure 6). In addition, in all three areas the correlations between the prediction accuracy and appearance measures were not significant (Figure 7). The analysis rejected the hypothesis that models were equiprobable (BOR < 0.05 for both edge and surface conditions), but the model for which the evidence was strongest was the same (M11) for both conditions (Edge: PEP = 0.039; Surface: PEP = 1.4e-4) (Supplementary Figure S5b).

### 4. Discussion

In the current study, we compared cortical activity related to color appearance generated by either edge-induced filling-in or a uniform surface chromaticity and found them to be associated with complementary patterns of activity in dorsal and ventral visual streams. Three lines of evidence support these results. First, the MVPA technique showed a significantly different profile of classification performance for the edge-dependent vs surface dependent conditions in dorsal areas V3A and V3B/KO compared to ventral areas V3v, hV4 and LO. These results are robust as these areas showed the same pattern of responses when replicated with a smaller stimulus and also when re-analyzed with ROIs that excluded the voxels retinotopically activated by the contours. It is unlikely that differential deployment of attentional mechanisms could explain these results as attention was uniformly controlled in all fMRI experiments. Second, individual differences in the psychophysically measured strength of the WCE were significantly correlated only with the classification performance of the edge-dependent stimulus in dorsal areas V3A and V3B/KO. Finally, a DCM analysis supported a model in which the edge-dependent stimulus modulated the effective connectivity directed from dorsal area V3A to area V1 and ventral area LO. In contrast, complementary modulations of LO onto V1 and V3A were found for the surface-dependent comparison with the control. Importantly, given that the whole brain GLM analyses failed to detect significant differences among the stimulus conditions, the results obtained depended critically on the sensitivity of the fine-grained MVPA (Sapountzis et al. 2010) and the DCM methods (Stephan et al. 2010, Friston, Harrison, and Penny 2003). Taken together, the results demonstrate that signals associated with edge- and surface-dependent information are not represented identically across visuals areas. The results support the hypothesis that information about surface-dependent and edge-induced colors is transmitted through distinct neural pathways.

The results do not rule out the participation of other areas in mediating edge- and surface-dependent color appearance. Indeed, the MVPA results reveal classification performance above chance in all visual areas, although they do not all significantly distinguish between the three conditions. For example, it would be interesting to extend the DCM analyses to include area V2 because of its implication in color processing (Roe and Ts’o 1999) and contour extraction (Lee and Nguyen 2001). This would entail examining a space of effective connections between four areas with an accompanying combinatorial increase in the number of possible models that would need to be tested. The larger number of models would dilute the power of such an analysis without, for example, evidence from the MVPA to constrain the analysis to a relevant subset of hypotheses.

### 4.1. Implication of cortical areas in edge- and surface dependent color processing

While area V3A has not typically been associated with color processing, activity dependent markers revealed a sparse population of cells responsive to chromatic stimuli (Tootell et al. 2004, Tootell and Nasr 2017). The small size of these color-selective cells and their random distribution may explain the failure to detect color-selective responses in this area (Brouwer and Heeger 2009). In addition, Hadjikhani et al. (1998) included V3A in a set of areas that generated greater responses to color than luminance modulated stimuli. More recently, Castaldi, et al. (2013), reported that V3A responds to chromatic spatial features. Its influence on color areas would be made possible by the dense connectivity between dorsal and ventral streams (Markov et al. 2013) made possible in human via the Vertical Occipital Fasciculus (Takemura et al. 2015).

Area V3A exhibits response selectivity to stimulus features that are characteristic of the WCE. V3A has been shown to be selective for contour curvature (Caplovitz and Tse 2007), a characteristic of the inducer contours that is necessary to enhance the WCE but not sufficient to generate it alone, e.g., as shown by the braided control contours here and in previous work (Devinck et al. 2014, Gerardin et al. 2014). Area V3A is also implicated in filling-in phenomena, in that it was reported to show increases in activity during Troxler fading (Mendola et al. 2006). Areas V3 and V4 have recently been implicated in form-contingent filling-in induced by closed contours (Hong and Tong 2017). Similarly in macaque, area V3 has been implicated in filling-in phenomena (De Weerd et al. 1995). V3A also shows much larger spatial summation than do nearby cortical areas (Press et al. 2001) as might be expected for an area involved in filling-in over extended regions of the visual field. In fact, the large areas of the visual field over which the WCE can extend require mechanisms that are effective in the peripheral visual field. The implication of dorsal areas in large scale filling-in is consistent with the bias of peripheral visual field projections in the dorsal stream (Kravitz et al. 2013).

Early studies that reported retinotopically distributed neural activity due to brightness and color filling-in in area V1 (Sasaki and Watanabe 2004) have been disputed (Zweig et al. 2015, Cornelissen et al. 2006). Using fMRI, Cornelissen et al. (2006) found that surround-induced responses in V1 did not depend on surround intensity but instead could be attributed to an extended edge-response. This is consistent with voltage-sensitive dye-imaging in macaque that shows only edge responses to uniform fields of color and luminance in V1 (Zweig et al. 2015). Instead, V1 responses to contours have been attributed to figural effects rather than brightness or color filling-in (Kok and de Lange 2014, Cornelissen et al. 2006, Lee and Nguyen 2001). Interestingly, the WCE stimulus is reported to demonstrate both a long-range color filling-in and a strong influence on figure-ground segregation (Von der Heydt and Pierson 2006, Pinna, Brelstaff, and Spillmann 2001). However, the results from area V1 are not likely to be explained uniquely by figure-ground processing. The classification results from this area (Figure 6) were based only on the voxels interior to those activated by the retinotopic positions of the contours (blue regions in Figure 5). The highest prediction accuracy for V1 significantly differed from that for both of the other classification performed and pitted the surface-dependent stimulus against the braided control, i.e., stimuli having the same braided contour. Instead, this supports that classification depended on the interior chromaticity and/or the perceived color and not local or long-range contour detection (Table 1). The poor correlation of the classification accuracies from this area with the perceived strength of the filling-in color (Figure 7) would support the former of these two interpretations.

Areas V3v, hV4 and LO displayed their highest classification accuracy to the surface chromaticity of the stimulus and not to the edge-induced fill-in conditions. In line with this outcome, the classification accuracies for these areas were not correlated with the strength of the perceived filling-in color. These results are consistent with previous studies that have demonstrated strong responses of these ventral stream areas to uniform surface colors (Brouwer and Heeger 2009, Parkes et al. 2009, Bouvier, Cardinal, and Engel 2008, Bartels and Zeki 2000, Hadjikhani et al. 1998, Sakai et al. 1995).

Area V2 presents a more complex pattern of results in that the classifiers trained for this area showed high accuracy for all three conditions with no selectivity between comparisons. This pattern of results is consistent with neurophysiological studies in primates that show diverse cell classes in area V2 including those responsive to uniform color fields (Peng and Van Essen 2005, Roe and Ts’o 1999) and others that might be involved in contour detection and color induction (Roe and Ts’o 1999).

### 4.2. Role of double- and single-opponent cells

The initial segregation of responses to uniform color fields and edges in neural sub-populations occurs as early as area V1 and may reflect the differential responses of single- and double-opponent cells. Population responses to uniform color fields in area V1 are dominated by edge responses as demonstrated by optical imaging with voltage sensitive dyes (Zweig et al. 2015). This result could reflect that double-opponent cells are the most numerous color sensitive cell class in area V1 at the eccentricity examined. The less numerous single-opponent cells would be expected to respond as well to uniform fields. Thus, the typical cortical response signature in area V1 to a uniform color field could be expected to be a pattern of activity across cells with both types of receptive field profile. Isolated chromatic edges and edge transients, however, would be expected to generate a response profile dominated by double opponent cells.

Why then do chromatic edge-transients, such as in the case of the WCE, suffice to generate the appearance of a weakly colored field, filling-in the interior of a uniform region interior to the chromatic contour? It is significant that the color filling-in requires continuity of the bi-chromatic contour as shown here and elsewhere (Devinck and Knoblauch 2012) by the lack of effect for the braided control stimulus. This suggests a mechanism by which the coherence along the contour is integrated across a population of double-opponent cells’ responses that we hypothesize would be mediated via paths from V1 to V3A. Such an interpretation is supported by results from adaptation studies demonstrating the role of contours in generating filling-in phenomenon (Coia and Crognale 2017, Hazenberg and van Lier 2013, van Lier, Vergeer, and Anstis 2009). We suppose then that the coherent contour would signal a uniform color field rather than just an edge-transient through the concurrent modulation of ventral color areas and V1 via dorsal stream areas implicated in contour integration.

### 4.3. Distributed nature of the responses

We find responses to the stimuli distributed across multiple streams and at multiple levels of the visual hierarchy, in line with an interareal network which is observed to be much denser than previously thought (Markov et al. 2013) and with evidence of the highly distributed nature of object processing (Konen and Kastner 2008). In area V1, the induced activity of voxels remote from the contours is unlikely to be mediated by monosynaptic lateral connections because intrinsic connections extend over 2 mm or less (Markov et al. 2011), and given a cortical magnification factor of 6-13 mm/deg (Van Essen, Newsome, and Maunsell 1984, Daniel and Whitteridge 1961), it is unlikely that lateral activity spreads much larger than a degree (Angelucci and Bressloff 2006). This suggests that feedback projections from higher order areas play a role in the activation of area V1. The involvement of descending pathways is highly relevant given the role of feedback pathways in contextual processing (Zipser, Lamme, and Schiller 1996). Here, feedback projections from areas V3A and V3B/KO could be particularly significant given that we show that these areas code the behavioral response and that the dynamic causal model that best supports the data indicates that the effective connectivity from V3A to V1 is modulated differentially by the WCE. Recent human studies suggest that area V3A is relatively low in the cortical hierarchy (Michalareas et al.2016) so that these findings are in line with previous evidence of activity at early hierarchical stages reflecting the content of consciousness (Leopold and Logothetis 1996). In the framework of predictive coding, that postulates that prediction errors are propagated up the cortical hierarchy and predictions down the hierarchy (Friston and Kiebel 2009), the descending prediction signal from area V3A might be related to figure-ground perception which is thought to be represented in area V1 (Kok and de Lange 2014, Cornelissen et al. 2006, Von der Heydt and Pierson 2006) or alternatively a prior indicating the presence of a uniform color field.

## 5. Conclusion

Our results provide evidence that the visual system employs separate networks for the processing of surface colors that are generated by a field of uniform chromaticity and those due to filling-in induced by distant chromatic edges, so that the neural responses to a uniform and a filled-in color are not isomorphic. Responses in early visual areas are likely due to differential feedback processes from the dorsal and ventral streams. We, thus, conclude that surface color representation depends on a context-sensitive network of multiple distributed processes in the cortex.

## Acknowledgements

The authors thank Elisabeth Baumgartner, Karl Gegenfurtner, Stewart Shipp and Andrew Coia for comments and criticisms on a previous version of this manuscript. They also thank Mathieu Ruiz for help with data analysis and the technical staff at the Grenoble MRI facility IRMaGe.

